# Empathy for pain persists across live two-way video interactions and viewing of prerecorded videos

**DOI:** 10.1101/2025.07.07.663421

**Authors:** Jannik Heimann, Pauline Petereit, Anat Perry, Ulrike M. Krämer

## Abstract

The impact of digitally mediated social interaction on understanding others and sharing their emotions has not been thoroughly investigated. We examined how live, video-mediated interaction, as opposed to watching a prerecorded video, affects behavioral, neural, and physiological aspects of empathy for pain. Thirty-five observers watched targets undergoing painful electric stimulation in an electroencephalogram study. We hypothesized that reduced temporal presence, the immediacy or delay with which information is transferred during social interactions, would result in diminished behavioral and electrophysiological empathic responses. However, observer’s behavioral empathic responses were not diminished with reduced temporal presence. On a neural level, mid-frontal theta was sensitive to the other’s pain intensity, and we observed significant physiological coupling between participants. Mu suppression, on the other hand, was not modulated by pain intensity. Importantly, neural and physiological indices of empathy were independent of temporal presence. However, exploratory analyses indicated a latency effect of temporal presence on pain-related theta activity with an earlier theta increase in interactions with high temporal presence. The results suggest that the temporal presence of individuals may not be necessary for empathy towards another’s pain. Future studies may investigate more naturalistic social interactions and include motivational aspects of empathy. We discuss implications of these findings for debates on social presence and on second-person neuroscience.

---

Digitally mediated social interactions have become an integral part of everyday life, from social media to video conferencing to online psychotherapy. However, the implications of mediated social interactions for our understanding of others remain poorly understood. We recently proposed a theoretical framework for studying the effects of social presence on social cognition, in particular on empathy and mental state attribution (Krämer et al., 2025). In this framework, we defined social presence as the degree to which others are physically and temporally present, that is the degree to which verbal and nonverbal cues about others are available (physical presence) and the immediacy with which these social cues are available (temporal presence). Extending a recent study of ours, in which we investigated the influence of physical presence (Petereit et al., 2023), we here addressed how temporal presence affects empathy for pain on behavioral, neural and physiological levels.

Temporal presence refers to the immediacy or delay with which information about the other is transferred during social interactions, which in turn allows for the possibility of reciprocity. For example, a direct conversation between two individuals demonstrates high temporal presence due to the instantaneous exchange of information, while observing a prerecorded video reflects low temporal presence because of the temporal delay between recording and viewing (Krämer et al., 2025). Temporal alignment in social situations can be assumed to play an essential role in facilitating social coordination and communication (Cañigueral et al., 2022; De Jaegher et al., 2010; Gallotti et al., 2017; Schilbach et al., 2013). Only with high temporal presence, interacting individuals can directly react to the other’s behavior creating feedback-loops of reciprocal social exchange (Dumas, 2011; Hamilton & Holler, 2023). Compared to live, bidirectional interactions, watching a video of another person reduces the temporal presence of that person, which could also impair empathy due to the absence of reciprocal social cue exchange. We understand empathy as an emotional response that involves affective sharing of another’s feelings and cognitive understanding of their mental and emotional states, while maintaining a distinction between one’s own and the other’s states (Cuff et al., 2016; Zaki & Ochsner, 2012).

The effects of temporal presence in social interactions have been investigated especially in regard to eye gaze as a social communication signal. The social significance and the signaling function of direct gaze are impaired when temporal presence between interacting individuals is prevented (Hamilton, 2016; Hamilton & Lind, 2016; Risko et al., 2016). When comparing direct, real-life interaction, video-mediated interaction, and watching a prerecorded video, the skin conductance response to direct gaze of others differed only between prerecorded videos and direct interaction, suggesting that the direct gaze in the condition of preserved temporal presence was the critical factor for increased autonomic arousal (Hietanen et al., 2020). Moreover, patterns of gaze behavior were altered when comparing direct, live interactions and prerecorded videos, indicating differences in implicit mentalizing between direct and mediated interaction (Morgan et al., 2022). Research on action observation in infants, adults, and in macaque monkeys suggests stronger neural motor responses in live, direct action observation compared to watching video-taped actions (Caggiano et al., 2011; Järveläinen et al., 2001; Karimova et al., 2024; Shimada & Hiraki, 2006). A recent study comparing live and video-taped theater performances reported more skin conductance responses when observing live performances, suggesting increased physiological arousal with higher temporal presence (Gabriel et al., 2025). In a further study, mimicry and the development of trust during interactions remained consistent across direct, face-to-face interactions and live, video calls, whereas these features were found to be diminished in prerecorded video interactions (Diana et al., 2023), which may indicate reduced experience sharing with reduced temporal presence. These studies showed that temporal presence affects important components of empathy and associated constructs, including implicit mentalizing, mimicry, arousal in social situations and action sharing. Based on this, we hypothesized that temporal presence also impacts more complex facets of empathy, which has not been shown before. Thus, the current study investigated how reduced temporal presence affects our empathic abilities to share experiences and understand others in pain on behavioral, neural and physiological levels (Cuff et al., 2016; Decety & Jackson, 2004; Lieberman, 2007; Shamay-Tsoory, 2011; Zaki & Ochsner, 2012).

As in our previous study on physical presence and pain empathy (Petereit et al., 2023), we used an established empathy-for-pain paradigm (Lamm et al., 2011; Rütgen et al., 2015; Singer et al., 2004; Singer & Lamm, 2009). In this paradigm, the behavioral readout, also called “empathic accuracy,” is defined as the relationship between the ratings of the pain observer and the pain receiver (the target) (Gauthier et al., 2008; Ickes et al., 1990; Jospe et al., 2020; Zaki & Ochsner, 2011). Empathic accuracy is considered to reflect the cognitive dimension of empathy (Mackes et al., 2018; Rütgen et al., 2015), whereas the observer’s subjective experience of unpleasantness in response to the target’s pain is thought to reflect the affective component of pain empathy (Mu et al., 2008; Rütgen et al., 2015).

The neural correlates of empathy for pain are often described within the framework of shared representations and simulation theories of empathy. These theories posit that observers represent the targets’ experience in a similar manner and in corresponding brain areas (Decety & Jackson, 2004; Keysers & Gazzola, 2006; Lamm et al., 2011; Marsh, 2018; Preston & De Waal, 2002), which are sometimes referred to as mirror neuron system (Rizzolatti & Craighero, 2004). Shared representations are assumed to facilitate social understanding, and this mechanism has been proposed as a neural substrate for experience sharing and affective components of empathy (Bekkali et al., 2021; Gallese et al., 2004; Keysers & Gazzola, 2006; Singer & Lamm, 2009), including empathy for pain (Hari & Kujala, 2009; Riecansky & Lamm, 2019). In the current study, reduced temporal presence disrupts the exchange of social cues and contingent responding, or mutual adjustment. This disruption may weaken engagement of neural circuits related to simulations and the subsequent empathic processes. Accordingly, we focused on specific neural and physiological indices of shared representations and pain empathy, namely the mu-rhythm, frontal theta and “physiological coupling”. The so-called mu rhythm (8-13 Hz) stems from the somatosensory cortex and has been repeatedly associated with pain empathy, with mu power being suppressed in response to the pain of others (Cheng et al., 2008; Peled-Avron et al., 2018; Peng et al., 2021; Perry et al., 2010; Riecansky & Lamm, 2019; Whitmarsh et al., 2011; Yang et al., 2009). Frontal theta activity (2-8 Hz; Misra et al., 2017; Mu et al., 2008; Peng et al., 2021; Petereit et al., 2023) possibly stems from the anterior and mid-cingulate cortex (ACC/MCC, Misra et al., 2017; Mitchell et al., 2008; Van Der Molen et al., 2017; Zhang et al., 2021) which is implicated in both first-hand and vicarious pain as part of the “pain matrix” (Cheng et al., 2007; Fallon et al., 2020; Hutchison et al., 1999; Jackson et al., 2005; Lamm et al., 2011; Xiao & Zhang, 2018), the core network for nociceptive processing (Legrain et al., 2011). This is evidenced by its response to images of hands in painful situations (Mu et al., 2008) and to facial expressions of varying pain intensities (Saarela et al., 2006). Images of hands and faces in pain have both been shown to be associated with behavioral and neural aspects of empathy for pain (Gallo et al., 2018; González-Roldan et al., 2011; Goubert et al., 2005; Jauniaux et al., 2019; Murata et al., 2020; Saarela et al., 2006). Emotional facial expressions play a major role in social signaling (Cañigueral et al., 2022; Güntekin & Başar, 2014). Reduced temporal presence may hamper this reciprocal signaling by limiting the ability to express compassion or support. In mediated interactions, the salience of another’s suffering can diminish due to missing reciprocity and reduced self-relevance of the other’s experience (Conway et al., 2016; Schmitz & Johnson, 2007), as meaningful reactions are constrained (Winczewski et al., 2016; Zaki, 2014). This decreased self-relevance may lower engagement with the other’s pain, leading us to expect reduced neural responses under reduced temporal presence.

Another physiological measure of (pain) empathy is “physiological coupling”, which refers to the phenomenon of synchronized autonomous activity patterns, such as those observed in electrodermal (EDA; skin conductance response, SCR) or cardiac activity (interbeat interval, IBI) across interacting humans (Neumann & Westbury, 2011; Palumbo et al., 2017). Several studies have demonstrated the relevance of physiological coupling in contexts of empathy (Jospe et al., 2020; Levenson & Gottman, 1983; Pan et al., 2023; Qaiser et al., 2023), as well as in the context of pain perception (Goldstein et al., 2017; Murata et al., 2020; Reddan et al., 2020), self-reported feelings of social presence (Järvelä et al., 2016), and the effects of physical presence on empathy for pain (Petereit et al., 2023). Investigating the relationship between physiological coupling and empathic accuracy has provided mixed results (Jospe et al., 2020; Levenson & Ruef, 1992; Thorson, 2018; Zerwas et al., 2021).

In our previous study, we investigated the effects of ***physical presence*** on empathy for pain using the above-mentioned pain empathy paradigm (Petereit et al., 2023). Participants observed another person undergoing painful electric stimulation, either in a direct, face-to-face or in a live, video-mediated interaction. Physical presence did not affect empathic accuracy and shared unpleasantness, and neither did it affect pain-related mu suppression. However, observer’s theta power to the other’s pain and the physiological coupling between participants were diminished in mediated compared to direct interactions. These results suggest that physical presence, i.e., the richness of social cues in an interaction, is important for a nuanced physiological resonance with the other’s pain experience.

Building on this work, we examined how **temporal presence** impacts pain empathy by comparing live video calls with watching prerecorded videos of others in pain. Since reductions in temporal presence disrupt the immediate exchange of information and reciprocity, we hypothesized that temporal presence would affect empathy-related neurophysiological mechanisms in a similar way to, or even more than, physical presence (Petereit et al., 2023). In detail, we hypothesized that reduced temporal presence would result in diminished empathic responses. Specifically, at the behavioral level, we expected reduced temporal presence to result in decreased empathic accuracy and less shared unpleasantness of other’s pain. At the physiological level, we hypothesized that heartbeat and skin conductance coupling would decrease with reduced temporal presence. Finally, at the neural level, we expected mu suppression and midfrontal theta activity to vary with levels of temporal presence.

## Methods

### Participants

Thirty-five students from the University of Lübeck were recruited (28 females, 7 males, self-identified; mean age: 24 years (*SD* 5.7)). All participants served as target and observer. Before the experiment, we made sure that the interacting participants were unfamiliar with each other. We used the same power analysis as Petereit et al. (2023) since one can expect that the effect sizes are comparable based on the similar experimental conditions. The power analysis showed that the sample size of *N*=30 is sufficient to detect a small effect (*f²*=0.02) of condition on empathic accuracy in a generalized linear mixed model on single trials with a power of at least 0.98. The power calculation was done in R using data simulation in the *simr* package (Green & MacLeod, 2016). Exclusion criteria included current and past neurological, current psychiatric or cardiovascular disorders, current or chronic pain disorders or current pain medication intake.

Due to the experimental design, the very first participant only served as target and the very last participant only served as observer (see Experimental Procedure for further explanation). One participant had to be excluded because of a non-reported sleeping disorder and could neither serve as observer nor as target. One participant’s behavioral data were lost due to technical issues. Some participants had to be excluded because of bad data quality and were excluded for each data analysis separately (EEG: 1, EDA: 1, own pain: 4 participants excluded). This led to the sample sizes of *N*=31 for the behavioral data, *N*=31 for the neural data in the observing conditions, *N*=31 for the SCR, *N*=32 for the ECG data, and *N*=30 for the neural data in the “own pain” condition. Participants received course credit or €12 per hour. All participants provided written informed consent before taking part in the experiment. The experiment was approved by the Ethics Committee of the University of Lübeck and conducted according to the Declaration of Helsinki.

### Experimental design

In prior social presence research, the terms “intimacy” and “immediacy” have been used for physical and temporal presence, respectively (Cui et al., 2013; Petereit et al., 2023; Short et al., 1976). We updated the terms to improve the precision and clarity of the two dimensions of social presence (Krämer et al., 2025).

The experiment had three conditions: own pain, observed pain via live video call, and observed pain via prerecorded video (Figure 1A). In the “own pain” condition, participants received electrical stimulation and rated their pain intensity on a visual analog scale (VAS). In the observed pain conditions, observers watched another participant (the target) receive the stimulation and rated both the target’s pain intensity and their own unpleasantness while watching, also using a VAS.

**Figure 1.**
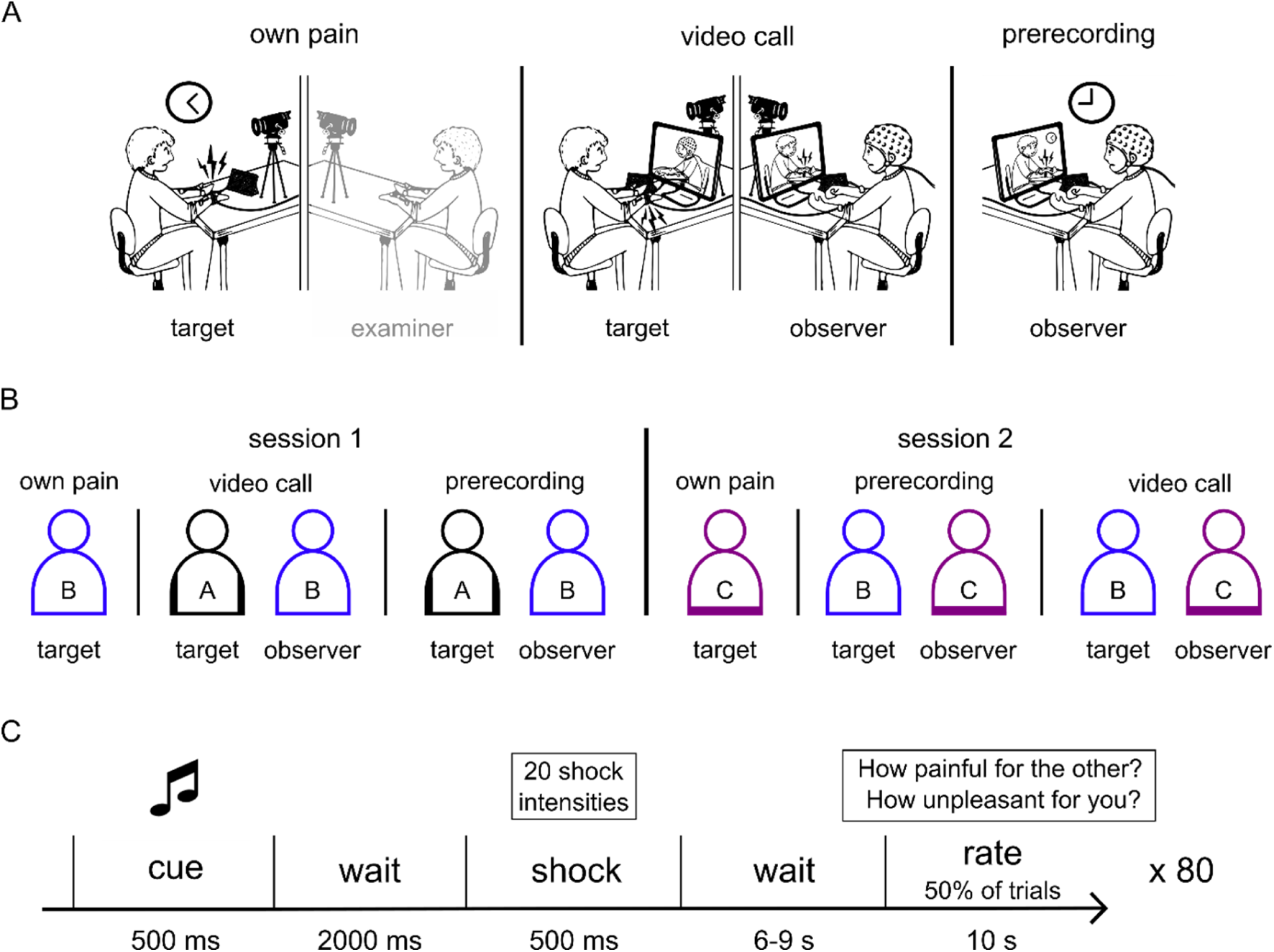
A) Schematic overview of experimental conditions. In the actual experiment, joysticks and not tablets were used for behavioral data acquisition and targets also wore EEG caps. B) Overview of the sequence of interactions between participants. Note that within session 1 participant B underwent the “own pain” condition first and then observed participant A. Participant B served as observer in session 1 and as target in session 2. The video that was recorded in the “own pain” condition in session 1 was presented in the “prerecording” condition in session 2. C) Overview of the trial structure with shock cue, shock and rating.

All participants completed all conditions and took on both roles (observer and target) across two laboratory sessions (Figure 1B). This within-subjects design differs from our earlier work (Petereit et al., 2023), in which participants served exclusively as either targets or observers.

The two primary conditions of interest were the counterbalanced live video call and prerecorded video conditions. In both conditions, observers viewed the target’s head, upper body, and the stimulated hand. In the “video call” condition, both participants could see each other in real time and were aware of the mutual visibility. In contrast, in the “prerecording” condition, observers viewed a recording of the target’s “own pain” condition, while the target could not see the observer. Cameras were positioned directly above the screens to approximate face-to-face interaction as closely as possible. The videos and the VAS were presented using PsychToolbox (PTB-3, Brainard, 1997; Kleiner et al., 2007).

To maintain comparability across conditions, the “own pain” condition was conducted via video call with an experimenter acting as observer and was recorded for later presentation in the “prerecording” condition.

Behavioral ratings, electrodermal activity (EDA), and electrocardiography (ECG) were recorded from both participants across all conditions. Electroencephalography (EEG) was recorded from observers during pain observation and from participants during their own pain experience, but not from targets while being observed. To ensure visual consistency, targets nevertheless wore EEG caps in all conditions. The experimental environment was standardized using gray standing walls surrounding the participants.

### Experimental procedure

Each participant attended two laboratory sessions. In the first session, participants completed the “own pain” condition and subsequently acted as observers in the “video call” and “prerecording” conditions, which were presented in randomized order. In the second session, participants performed the role of the target in the “video call” condition (Figure 1B).

After providing informed consent, participants read the instructions and were fitted with ECG, EDA, and EEG electrodes. This was followed by the pain stimulus calibration (see next section) and ten practice rating trials. Participants then completed the pain task in the “own pain” condition, followed by the two observing conditions. After each condition, participants answered questions assessing the perceived quality of the interaction. At the end of the experiment, they completed demographic and personality questionnaires.

#### Pain stimulus calibration

Before the actual experiment, the pain stimulation was calibrated to the subjective pain thresholds of the participants. The calibration started at 0 mA shock intensity and increased by steps of 0.5 mA. The maximum possible pain stimulation was at 10 mA. Only for the calibration, participants rated each shock on a 9-point Likert scale from 0 (”not perceivable”) to 8 (”worst possible pain”). The in-between steps on the scale were labeled as 1 = “noticeable”, 2 = “unpleasant”, 3 = “slightly painful”, 4 = “medium painful”, 5 = “very painful”, 6 = “extremely painful, 7 = “unbearably painful”, 8 = “worst possible pain”. The shock intensity was increased until the participant rated the stimulus as 7 = “unbearably painful”. Then, the intensity decreased by steps of 0.5 mA until 0 mA or “not perceivable” was rated. This procedure of increasing shock intensity was then repeated once. The final stimulus intensity which was rated as 1 = “noticeable” was used as the lower limit and the intensity rated as 6 = “extremely painful” was used as the upper limit in the actual experiment. Pain stimulus calibration results did not differ between the “own pain” and “video call” conditions (lower limits: own pain *M* = 0.9 mA, video call *M* = 0.9 mA, *t*(33) = 1.22, *p* = .231; upper limit: own pain *M* = 6.5 mA, video call *M* = 6.6 mA, *t*(33) = 0.38, *p* = .703).

#### Pain stimulation task

The pain stimulation task was very similar to that described by Rütgen et al. (2015). A Digitimer DS 5 isolated bipolar constant current simulator (Sprenger et al., 2020) was used with a Digitimer bar electrode (9 mm diameter, 30 mm apart, 10 volts) attached to the back of the left hand. Unlike in our preceding study, we attached the electrode to the left and not the right hand to facilitate handling of the right-hand joystick for the rating task. We used an abrasive gel underneath the electrode to improve the electric conductivity of the skin. A schematic overview of a trial is given in Figure 1C. The pain stimulation followed an auditory 500-ms cue, which was used for all different pain intensities alike. Between the cue and the stimulation was a 2000-ms interval in all trials. The electric shock itself was delivered for 500 ms in a series of 2-ms electric pulses, interspersed with 20-ms breaks. There was an inter-trial interval with a pseudo-randomized duration of 6-9 seconds. The pain intensity ratings were given in a total of 50% of the trials and were initialized with a vocal recording saying “Please rate”. The VAS ranged from 0 = “not painful at all” to 100 = “extremely painful”. Targets rated the pain intensity for themselves and observers rated the perceived targets’ pain experience. Further, observers rated how unpleasant it was for themselves to witness the others’ pain experience on a VAS from 0 = “not unpleasant at all” to 100 = “extremely unpleasant”.

The range of the electric shocks were evenly split into twenty different levels of intensities in the range obtained from the pain stimulus calibration for each participant. In total, there were 80 trials with each intensity appearing four times in each of the three experimental conditions. This is the same number of trials used previously in empathy for pain tasks (Petereit et al., 2023; Rütgen et al., 2015), and rating 50% of the trials is sufficient according to prior power analysis. Shock intensity ordering was fixed in all conditions and for all participants and was pseudo-randomized with less than four low- or high-intensity shocks in a row. Targets were instructed to express their pain experience as they wished and the observers were instructed to infer the others’ pain as accurately as possible.

### Questionnaires

Following the completion of each experimental condition, participants were prompted to indicate how immediate they experienced the interaction, and how close towards and how attached they felt to the other participant. These measures of perceived social presence were used to validate the effectiveness of the social presence manipulation and were tested for differences between conditions using a paired *t*-test. See Supplementary materials S1 for information about additional questionnaires that were not analyzed in the current study.

### Physiological data acquisition

EEG data were recorded with 64 sintered Ag/AgCl sensors in a cap with integrated eye and ear lobe electrodes. Electrode placement was according to the international 10-20 system. The online reference was placed on the right earlobe and an offline reference was on the left earlobe. Horizontal and vertical EOG were recorded with 4 electrodes, including Fp2 as the upper vertical electrode, one lower right vertical electrode and two horizontal electrodes next to the outer corners of the eyes. Data were recorded with a Brain Products GmbH system (BrainAmp MR plus amplifiers, BrainVisionRecorder software version 1.21.0102). The sampling rate was 500 Hz with an online high-pass filter of 0.016 Hz, low-pass filter of 48 Hz and a notch filter at 50 Hz. Impedances were kept below 5 kΩ.

ECG data were recorded with bipolar Ag/AgCl recording electrodes and one reference electrode, using a 50-Hz notch filter. One of the recording electrodes was placed on the left forearm, the other one on the right lower calf of the participant (following Einthoven’s lead II configuration, Quigley et al., 2024). Skin conductance was measured with 2 electrodes placed on the thenar and hypothenar of the right hand, using a 50-Hz notch filter. An electrode attached to the right forearm served as ground for both ECG and skin conductance, which were recorded with the same amplifier (BrainAmp ExG, BrainProducts GmbH). In the “video call” condition, data from the observer and target were recorded synchronously by connecting all amplifiers to the same USB adapter feeding the data into BrainVisionRecorder.

### Physiological data processing

#### EEG data

The EEG preprocessing was done using EEGLAB (version 2023.0, Delorme & Makeig, 2004) and ERPLAB (version 9.0, Lopez-Calderon & Luck, 2014), implemented in MATLAB R2022b (The Mathworks, 2022). First, data were re-referenced to linked earlobes and bipolar horizontal and vertical EOG channels were calculated. Noisy or bad data quality channels were marked to be ignored in further analysis (observing conditions: *M* = 3.1 channels, *SD* = 2.7; “own pain” condition: *M* = 7.1, *SD* = 2.8). Due to the first-hand experience of pain and consequent flinching, the data in the “own pain” condition was more artifactual than in the observing conditions. However, the affected channels were positioned in the peripheral areas of the skull, away from the central regions of the head that were of interest in the current study. In the next step, bad data segments or big artifacts were identified visually and removed. Before the independent components analysis (*runica* function in EEGLAB), data were bandpass filtered (non-causal Butterworth impulse response function in ERPLAB; lower cut-off: 1 Hz, upper cut-off: 40 Hz, 48 dB/oct roll-off). The ICA weights were back-projected to the original unfiltered data without removed bad data segments (Stropahl et al., 2018). This procedure guarantees accurate counting of rejected trials in the final step of threshold artifact rejection. Another bandpass filter was applied (non-causal Butterworth impulse response function in ERPLAB; lower cut-off: 0.2 Hz, upper cut-off: 40 Hz, 48 dB/oct roll-off) to the original data with the back-projected ICA weights. ICA components clearly related to ocular artifacts (visual inspection) were removed (observing conditions: *M* = 2.5 components, *SD* = 0.7; “own pain” condition: *M* = 2.7, *SD* = 0.8; Jung et al., 2000). Noisy channels were interpolated (spherical interpolation). Afterwards, data were epoched into segments time-locked to the shock onset with epochs ranging from -2500 to 5000 ms and a baseline correction from -1000 to 0 ms. Finally, a semi-automatized voltage threshold-based artifact correction was performed (−60/60 to -100/100 μV; rejected trials in observing conditions: *M* = 14%, *SD* = 3%, in “own pain” condition: *M* = 15%, *SD* = 4%).

In the “prerecording” condition, EEG triggers had to be sent based on what was happening in the video. Triggers in this case were sent using the logfile data from the corresponding previous participant’s “own pain” condition. However, since the video and the VAS for the observer had to be presented in parallel, trigger timing deviated slightly more and more variable than expected based on piloting. Because of this, trigger times in the “prerecording” condition were corrected based on the trigger times in the logfiles of the corresponding “own pain” condition when the video had been recorded. The correction of the trigger timings was performed based on the “experiment start” trigger as time point zero in the “own pain” condition, in which the video was recorded. This “experiment start” trigger corresponded to the start of the video in the “prerecording” condition and trigger timings were corrected according to this time point zero.

Finally, we conducted a time-frequency analysis according to recommendations from Cohen (2014). Data of all channels and in the frequency range of 1-40 Hz were Morlet wavelet convoluted (cycles evenly spaced from 5 to 14 cycles for the range of 1 to 40 Hz) after Laplace transformation (Cohen, 2014; Perrin et al., 1989). Power values were converted to decibels with a single-trial baseline normalization of -700 to -200 ms before stimulus onset. The time window of -500 to 2500 ms was extracted and downsampled to 100 Hz.

#### ECG data

ECG data preprocessing was done in MATLAB and EEGLAB. The data were bandpass filtered (non-causal Butterworth impulse response function in ERPLAB; lower cut-off: 1 Hz, upper cut-off: 30 Hz, 48 dB/oct roll-off) and segmented into 10 s epochs (3 s before and 7 s after stimulus onset). The R-peaks were detected using the MATLAB function *findpeaks*. Afterwards, the epochs were visually screened for wrongly detected or missing R-peaks. Extrasystoles and other artifacts were removed. The interbeat interval (IBI) in ms was calculated for every pair of heartbeats in each epoch. The IBI values were assigned to every data point in the epoch with the original sampling rate. Every epoch was baseline corrected to the mean baseline averaged across the two seconds before shock onset of all trials within each participant per condition. Outliers, defined as values exceeding ±3 standard deviation within each participant, were excluded before averaging, resulting in 18.2% excluded trials.

#### Skin conductance data

Skin conductance responses (SCR) were calculated using the Ledalab-Toolbox for MATLAB (Benedek & Kaernbach, 2010a, 2010b, Version 3.4.8). Data were downsampled to 50 Hz and smoothed with an adaptive Gaussian filter. Visually identified artifacts were spline-interpolated. To separate tonic and phasic electrodermal activity, a continuous deconvolution analysis was conducted (Benedek & Kaernbach, 2010b). For SCR amplitude analysis, SCRs greater than 0.01 µS were kept for further analysis (Boucsein et al., 2012). An additional artifact correction was performed before averaging trials, using a ±3 standard deviation exclusion criterion within each participant. Because participants operated the joystick with the same hand where the EDA electrodes were attached, artifacts sometimes occurred beyond the ITI during the first seconds of the next trial. Therefore, the correction was done for a time window of -1 to 1 sec around the shock onset which was before the onset of the SCR. The artifact correction resulted in 24.86% excluded trials. Individual SCR data were range-corrected by each participant’s maximum SCR (Hein et al., 2011; Lykken, 1972).

### Statistical analyses

Before the details of each analysis are described, we will give an overview of the general statistical analysis rationale:

Linear mixed models (LMM) were calculated for behavioral data, EEG power, and IBI data using R (R Core Team, 2020). A generalized LMM (GLMM) with a gamma family and log link was calculated for the right-skewed SCR data (skewness = 2.48). The fixed effects structure was dependent on the hypotheses given the type of analysis. The details are given in the following subsections. The model specifications for each final model are given in the Supplementary materials S2. The experimental condition (“video call” vs. “prerecording”) was dummy-coded by using the simple effect-coding scheme (Daly et al., 2016). The random effects structure was identified by starting with the most complex random effects structure (random intercept and random slope for condition*predictor-of-interest interaction for each observer) (Barr et al., 2013) and successively removing the highest-order random effects with the lowest variance until the model converged, as suggested by Singmann and Kellen (2019). SCR data required a Gamma log link function for positive, right-skewed data (Lo & Andrews, 2015). Accordingly, models were calculated with *lmer* and *glmer* functions in the *lme4* package in R (Bates et al., 2015). LMM significances were tested with the *lmerTest* package (Kuznetsova et al., 2017). The predictors target rating, shock intensity, physiological data and EEG power were z-standardized to guarantee (G)LMM convergence. *R²* marginal and conditional were calculated using the *MuMIn* package (Bartoń, 2010).

If the target’s data (independent variable) significantly affected the observer’s data (dependent variable), it would suggest that the target’s response influenced the observer’s response. Therefore, it would be considered empathic. A significant effect of condition (“video call” vs. “prerecording”) would imply that temporal presence influences the observer’s response. A significant interaction between target’s data and temporal presence would mean that the observer’s empathic response to the target’s pain response differed with the level of temporal presence. Depending on the direction of the effect, this could indicate increased or diminished empathic responses in the “video call” or “prerecording” condition. Thus, it would indicate increased or reduced empathy with lower or higher temporal presence. In the case of non-significant results for the (G)LLMs, we calculated Bayes Factors (Schmalz et al., 2023) to report evidence for or against the null hypothesis. Therefore, we used an approximation of the Bayes Factor that is calculated by using the BIC from the obtained (G)LMMs (Faulkenberry, 2018; Raftery, 1995). Since all mixed models test separate hypotheses, they do not require correction for multiple comparisons. However, all post hoc tests and correlational analyses were Bonferroni-corrected.

#### Behavioral data

The questions regarding the subjective quality of the interaction were compared between conditions with a paired *t*-test. For the pain rating data, analyses of empathic accuracy (based on perceived pain intensity) and of affective empathy (based on unpleasantness ratings) were conducted. The distinction between a cognitive (empathic accuracy) and an affective (shared unpleasantness) behavioral readout of empathy was based on classic definitions of empathy and prior studies showing differences in neural and behavioral correlates of these constructs (Cuff et al., 2016; Rütgen et al., 2015; Zaki & Ochsner, 2012). We related the observers’ ratings of the targets’ pain experience to the targets’ pain ratings as measure for empathic accuracy (Gauthier et al., 2008; Ickes, 1993; Ruben & Hall, 2013; Sun et al., 2018). We related the observers’ unpleasantness ratings to the targets’ pain ratings as measure for affective empathy (Rütgen et al., 2015; Shamay-Tsoory, 2011; Zaki & Ochsner, 2012). In the LMMs, we assessed whether observer’s pain and unpleasantness ratings were predicted by the interaction of the target’s pain ratings and the temporal presence condition.

Although we considered the observers’ ratings of the other’s pain and their own unpleasantness as indices of two different psychological constructs (empathic accuracy and affective empathy, respectively), we checked if these ratings were correlated. We calculated Pearson correlation coefficients for each dyad per condition separately. To average the resulting *p*-values of all correlations, combined p-values were calculated using Stouffer’s *Z*-score method for dependent *p*-values (Stouffer et al., 1949). To obtain the distance between observer and target ratings, we calculated the difference between these ratings for each dyad per condition separately and averaged the differences for each condition.

#### EEG data

First, we explain the analyses for EEG data according to our hypotheses based on the literature and based on the results of the whole time-frequency-electrode space analysis of Petereit et al. (2023). Second, we explain the analysis of the whole time-frequency-electrode space, which closely matches the analysis Petereit et al. (2023) presented. Finally, we explain the analysis of the own pain EEG data. For all statistical analyses of the EEG data, the calculated LMMs contained a fixed effects structure according to our hypotheses, including the interaction of temporal presence condition and the shock intensity. The mean power in each time-frequency cluster was calculated for each trial to be predicted by the model.

First, we analyzed the EEG data based on hypotheses derived from the literature for mu suppression and from our previous study on physical presence (Petereit et al., 2023) for theta and beta activity. For mu suppression, we analyzed the power within the frequency range 8-12 Hz at electrode C3 over the somatosensory cortex in a time window from 500 to 1000 ms after shock onset (Genzer et al., 2022; Hoenen et al., 2015; Perry et al., 2010). For theta activity, we looked at electrode Cz in a time-window of 100-600 ms at frequencies of 3-9 Hz, and for (low) beta activity, we looked at electrode CP1 in a time-window of 100-600 ms at frequencies of 11-20 Hz (Petereit et al., 2023).

Similar to our previous study, we conducted an exploratory whole-brain analysis by investigating the whole time-frequency-electrode space (1-40 Hz, all electrodes). The data was averaged across participants and permutation tests (*n*=1000) were calculated to obtain null hypothesis distributions (Cohen, 2014). Three different permutation tests were calculated to investigate the effects of shock intensity, condition and the interaction of both. For the effect of shock intensity, Spearman correlation coefficients of shock intensity and EEG power were calculated. In the permutation test, shock intensity labels were shuffled and randomly assigned. For the effect of condition, condition labels were shuffled and randomly assigned. Then, condition differences were calculated. For the effect of the interaction of shock intensity and condition, Spearman correlation coefficients of shock intensity and EEG power were calculated, shuffled and condition labels were randomly assigned. Then, condition differences were calculated. We used maximum value correction to correct for multiple comparisons. To identify the largest clusters and the location of the maximum activity that survived the correction for multiple comparisons, cluster sizes were calculated using the *bwconncomp* command in MATLAB. By this, the frequency boundaries and time window boundaries as well as the electrodes of maximum activity of the largest clusters were identified.

#### IBI and SCR data

To test for SCR coupling, observers’ SCR was predicted in a GLMM using a fixed effects structure of temporal presence condition in interaction with targets’ SCR. The physiological coupling was obtained by the relation of observers’ SCR with targets’ SCR. For the targets’ SCR, we considered the mean phasic activity in the time window of 2140-5140 ms after shock onset, which corresponded to the 3-second time window of the maximum averaged response of targets to the shock onset. For observers’ SCR, four time windows (2.5-5.5, 3-6, 3.5-6.5, 4-7 s) were tested to account for a possible time lag in observers’ responses to the targets’ pain expression. The 4-7 s time window was used for analysis, because it yielded the highest correlation of target’s SCR with observer’s SCR averaged across both conditions.

To test for IBI coupling, the same procedure as for SCR coupling was used, but with the IBIs as inputs for the LMM. For the LMMs, the mean IBI in the time window of 2136-5136 ms after shock onset was used for targets’ IBI. This was the 3-second time window (Sperl et al., 2016) of maximum averaged response of targets’ IBI to the shock onset. For observers’ IBI, five time windows (2.2-5.2, 2.5-5.5, 3-6, 3.5-6.5, 4-6.8 s) were tested to account for a possible time lag in observers’ responses to the targets’ pain expression. The 2.5-5.5 s time window was used for analysis, because it yielded the highest correlation of targets’ IBI with observers’ IBI averaged across both conditions.

### Exploratory analysis of the temporal presence effect on EEG latencies

The exploratory whole-brain analysis yielded effects for mid-frontal theta and right posterior alpha activity. During data analysis, we observed a latency difference between temporal presence conditions in these frequency bands. To further explore this, we calculated the peak latency for every participant in both observing conditions separately. Therefore, we automatically identified the maximum activity (Kompus et al., 2015) in the time window of 100 to 1500 ms after shock onset for theta (1-6 Hz, Cz) and alpha bands (9-13 Hz, PO8). The frequency bands resulted from the whole-brain analysis. The latencies of the peaks were compared between conditions using a paired *t*-test.

### Relationships between different empathy measures

We correlated different empathy measures to explore their relationship. Therefore, we calculated Pearson’s correlations between single-trial theta activation and pain or unpleasantness ratings. Secondly, we correlated empathic accuracy (correlation of observer and target ratings) and physiological coupling (correlation between observer and target IBI or SCR). Correlation coefficients were Fisher-*z* transformed before averaging across subjects. For comparisons across conditions, correlation coefficients were calculated per dyad and condition and then averaged. Combined p-values were calculated using Stouffer’s *Z*-score method for dependent *p*-values (Stouffer et al., 1949).

## Results

### Difference in perceived qualities of the interaction between temporal presence conditions

After each condition, we asked the observers how they perceived the quality of the interaction, specifically their feelings of immediacy, closeness of the interaction, and attachment to the other participant. Observers rated the immediacy (*t*(29) = 5.16, *p* < .001, *p_Bonferroni_* < .001, *d* = 0.94) and closeness (*t*(29) = 4.02, *p* < .001, *p_Bonferroni_* = .001, *d* = 0.72) higher after the video call compared to the prerecorded video, but did not rate their feelings of attachment differently (*t*(29) = 1.47, *p* = .153, *p_Bonferroni_* = .458, *d* = 0.28). Results are shown in Figure 2A. These results provide support for the construct validity of our paradigm.

**Figure 2.**
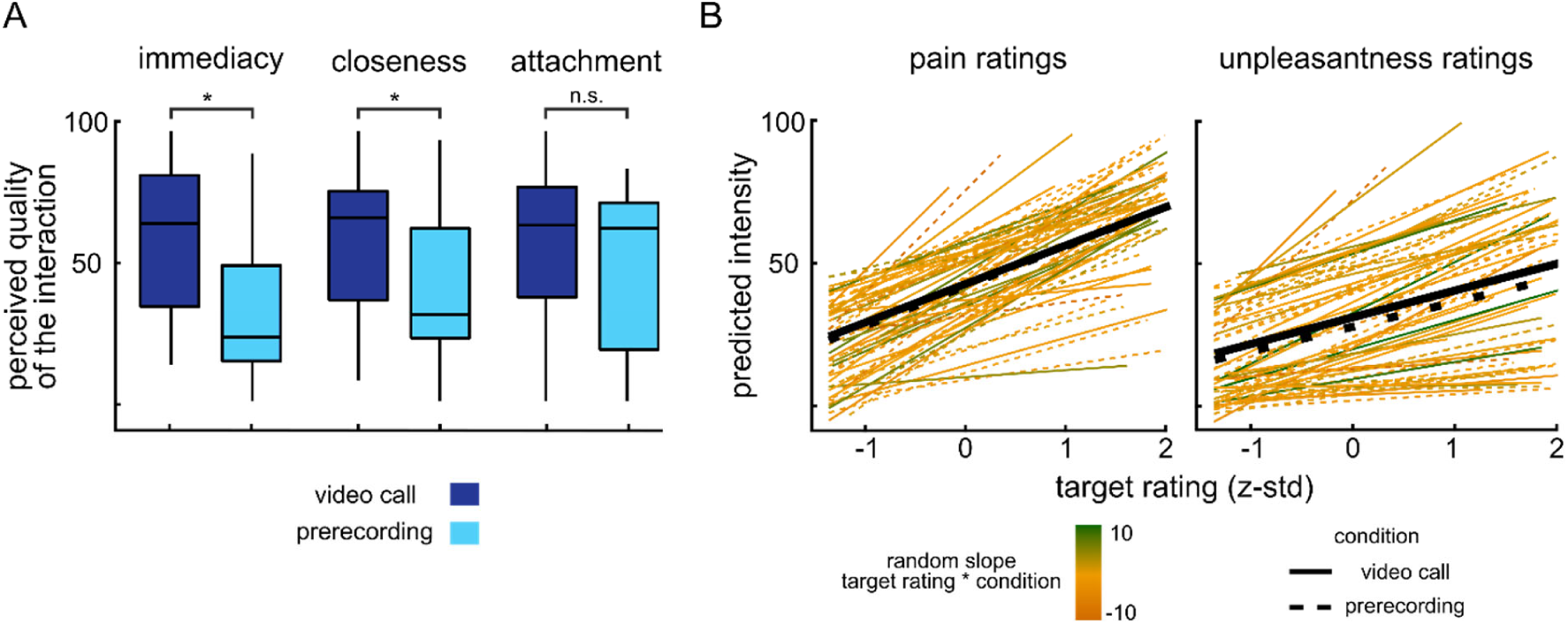
A) Observers’ perceived qualities of the interaction. After both conditions, observers were asked to rate the immediacy and closeness of the interaction with the target, and to indicate their attachment to the target. Condition differences were found for perceived immediacy and closeness but not for attachment. B) LMMs on single-trial behavioral data. Target ratings significantly predicted observers’ pain and unpleasantness ratings, which indicates empathic responses in observers. But this was similar in both conditions. Black solid and dotted lines represent the “video call” and “prerecording” averages, respectively, while the green to yellow color coding reflects the strength of random slopes per participant. * = *p*<0.001, n.s. = not significant.

### Temporal presence does not influence empathic accuracy

To assess empathic accuracy, we calculated a LMM that contained targets’ pain ratings, condition (“video call” vs. “prerecording”) and their interaction to predict observers’ pain ratings. Model parameters can be found in Table 1. Results show that observers’ pain ratings were predicted by targets’ pain ratings indicating meaningful empathic accuracy of the observers. However, this effect was not influenced by condition indicating that empathic accuracy did not differ between video calls and prerecorded videos (Figure 2B). Correlating observer and target pain ratings yielded similar coefficients for both conditions (video call: *r*(31) = .65, *p* < .001, *p_Bonferroni_* < .001; prerecording: *r*(31) = 0.64, *p* < .001, *p_Bonferroni_* < .001). The mean distance between observer and target pain ratings was 1.99 (*SD* = 15.8) in the “video call” and -1.57 (*SD* = 16.5) in the “prerecording” condition, indicating high accuracy in both conditions considering the 0-100 VAS scale. Testing the model with and without the fixed effect of condition, the obtained Bayes factor of BF_10_ = 0.001 indicates extreme evidence for the model without the predictor “condition” given our data. The results demonstrate that the observers’ empathic accuracy was equally high across levels of temporal presence.

**Table 1.**
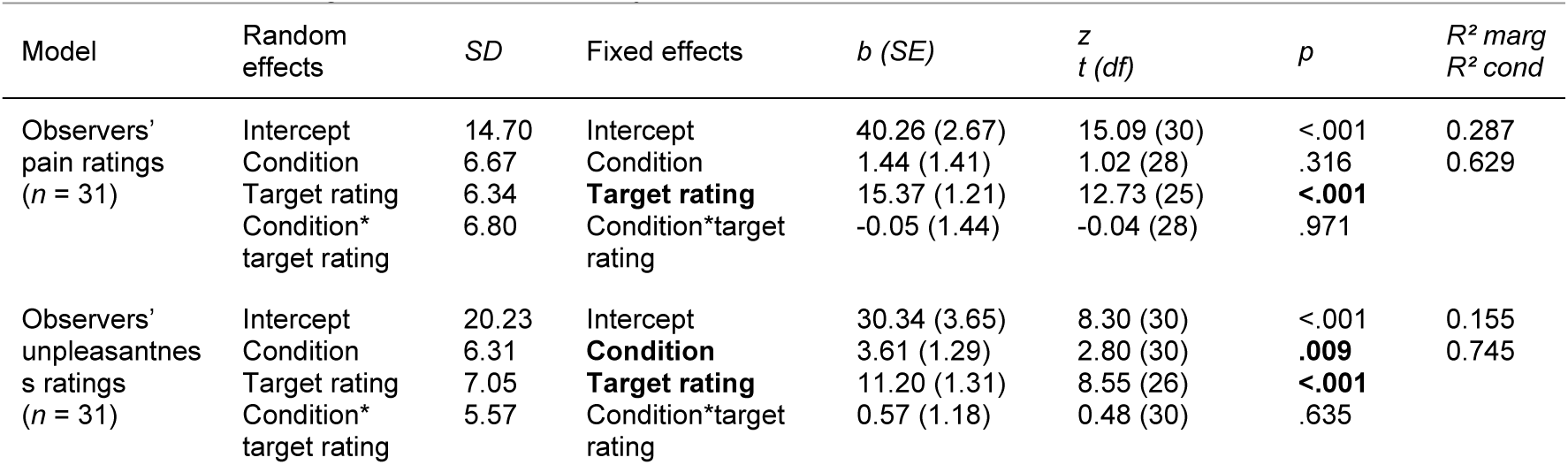
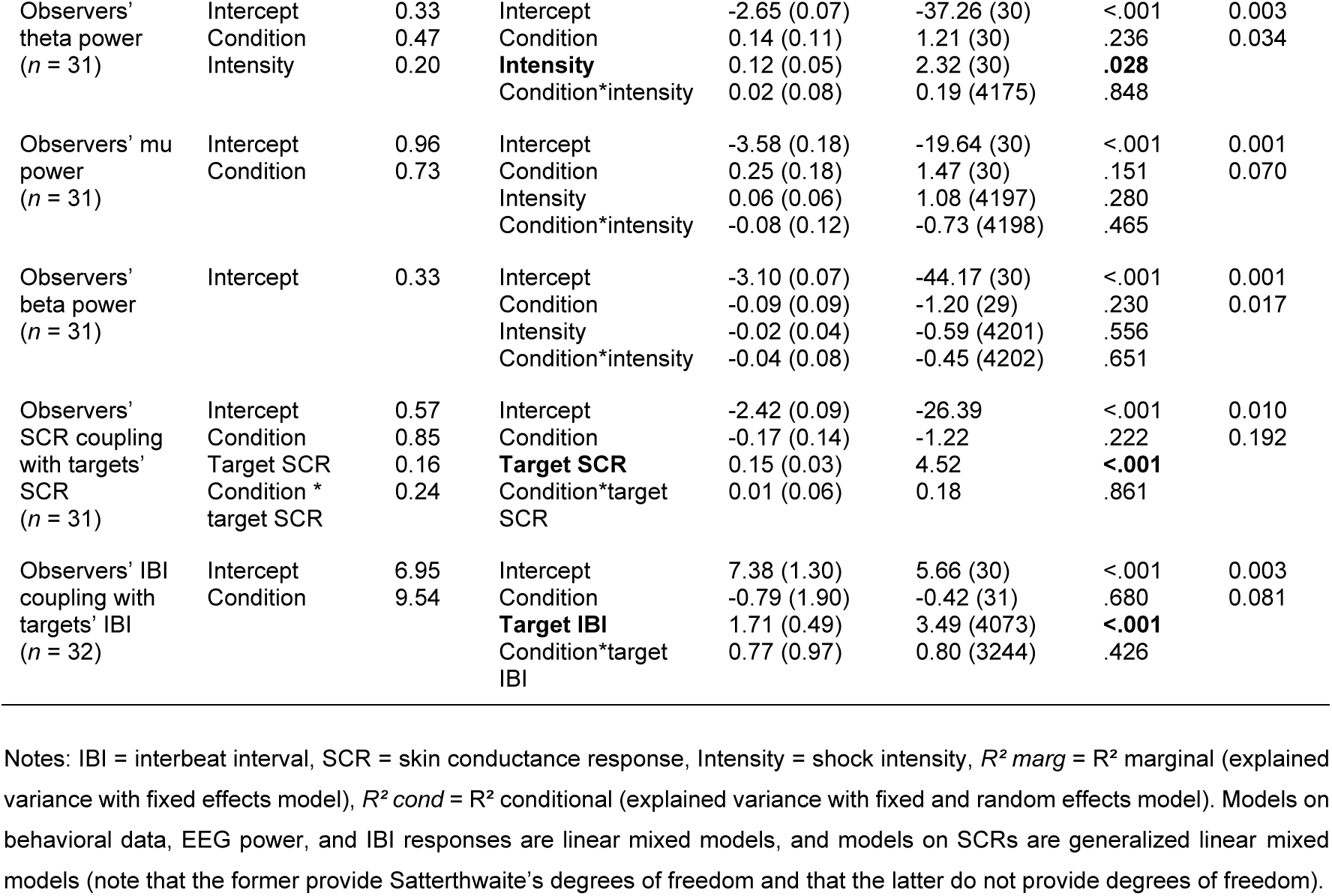
Results of the single-trial (generalized) linear mixed models on observers’ pain and unpleasantness ratings, affective empathy, IBIs, SCRs, and EEG theta, mu, and beta power.

### Temporal presence does not influence affective empathy

To assess affective empathy, we calculated a LMM that contained targets’ pain ratings, condition and their interaction to predict observers’ unpleasantness ratings. Model parameters can be found in Table 1. Results show that observers’ unpleasantness ratings were predicted by targets’ pain ratings indicating affective empathy of the observers. Further, observers’ unpleasantness ratings were generally slightly larger in the “video call” compared to the “prerecording” condition, indicated by a main effect of condition. However, the mean difference of only 1.66 (*SD* = 21) along the VAS, which ranged from 0-100, is negligible. Furthermore, the factors did not interact, showing that the effect of targets’ pain ratings on observers’ unpleasantness ratings did not vary with the temporal presence condition (Figure 2B). Correlating observers’ unpleasantness and targets’ pain ratings yielded similar coefficients for both conditions (video call: *r*(31) = .58, *p* < .001, *p_Bonferroni_* < .001; prerecording: *r*(31) = .57, *p* < .001, *p_Bonferroni_* < .001). Testing the model with and without the fixed effect of condition, the obtained Bayes factor of BF_10_ = 0.028 indicates very strong evidence for the model without the predictor condition given our data. The results indicate that the observers’ affective empathy was equally high across levels of temporal presence.

Although we considered the observers’ ratings of the other’s pain and their own unpleasantness as indices of two different psychological constructs (empathic accuracy and affective empathy, respectively), it should be noted that the two ratings were correlated. When calculating the correlation of ratings across all participants and for each condition, the results confirmed that there is a significant relationship between observers’ pain and unpleasantness ratings (video call: *r*(31) = .90, *p* < .001, *p_Bonferroni_* < .001; prerecording: *r*(31) = .90, *p* < .001, *p_Bonferroni_* < .001).

### Mu suppression, theta, and beta power are not sensitive to temporal presence

Based on the whole-brain analysis of our last study on physical presence (Petereit et al., 2023) and on the literature, we hypothesized that observers’ mu, theta and beta activity should be modulated both by the targets’ pain level and by the level of temporal presence. We tested these hypotheses by using LMMs on EEG power predicted by the interaction of condition and shock intensity in the frequency bands of theta (3-9 Hz, electrode Cz, time window 100-600 ms), mu (8-12 Hz, electrode C3, time window 500-1000 ms), and beta power (11-20 Hz, electrode CP1, time window 100-600 ms). Visualizations are shown in the middle and right column of Figure 3A-C. LMM model parameters can be found in Table 1.

**Figure 3.**
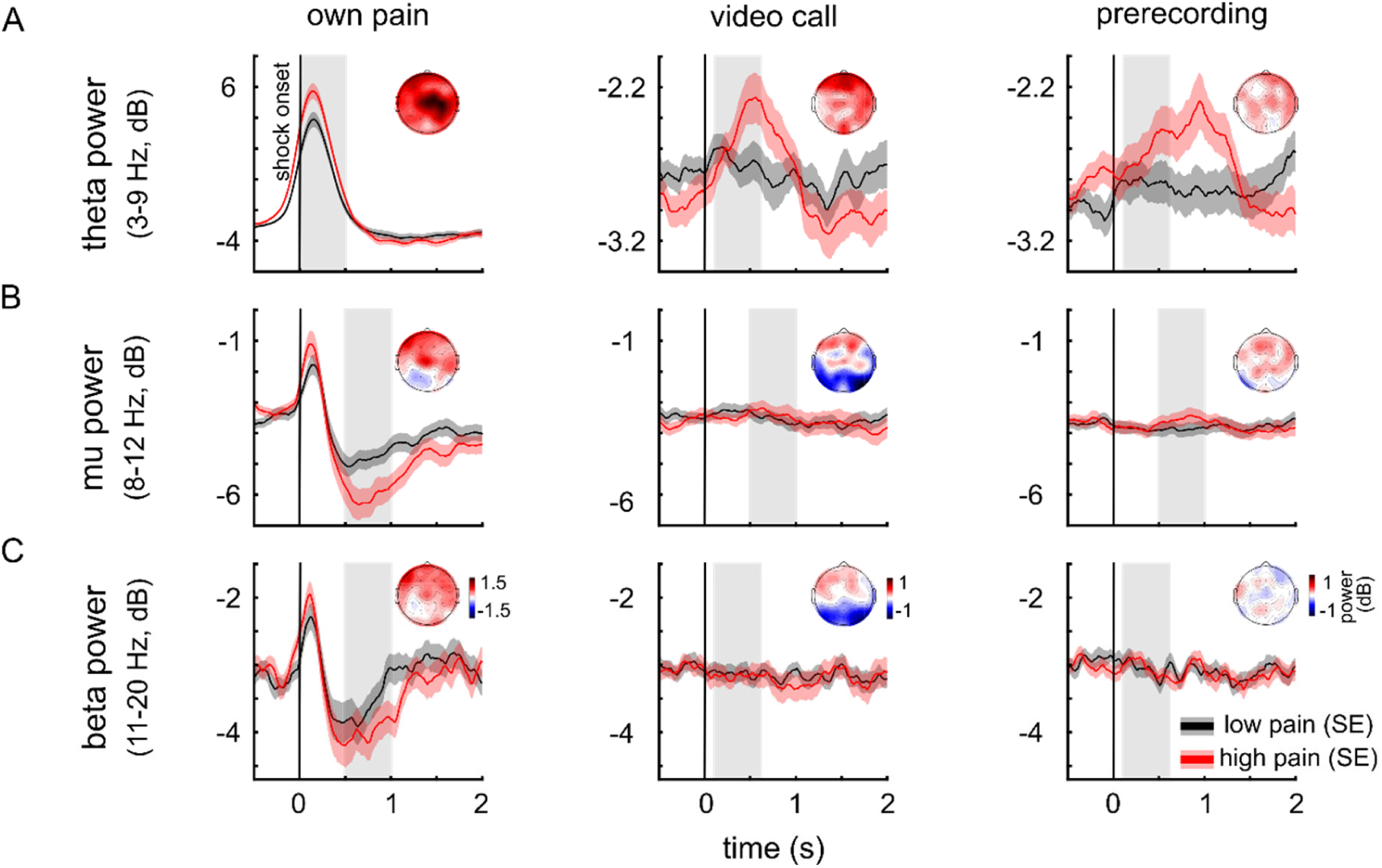
Time-power plots for the “own pain”, “video call”, and “prerecording” condition; for A) Theta (electrode Cz), B) mu (C3), and C) beta (CP1) oscillations. Participants’ own pain was related to theta, mu and beta activity. In the observing conditions, only theta activity was related to the shock intensity (Supplementary materials S3). This relationship did not differ significantly between the “video call” and the “prerecording” condition. Grey areas indicate hypothesized time windows for a shock intensity and condition interaction. Observers’ theta responses occurred later than hypothesized. Shock intensities were dichotomized into low- and high-intensity for display purposes only.

We found an effect of the shock intensity on observers’ frontal theta activity, indicating increased theta activity in observers in response to more intense pain in others. Shock intensity did not have an effect on observers’ mu and beta power. Importantly, there were no significant effects of condition or of the interaction of shock intensity and condition on mu, theta or beta activity. This demonstrates that temporal presence did not influence neural activity in the hypothesized frequency bands derived from the whole-brain analyses from our previous study. The Bayes factors for testing the effect of condition in the LMMs for theta, mu and beta activity confirmed that there was no temporal presence effect on EEG power in the given models (theta BF_10_ = 0.0005, mu BF_10_ = 0.0009, beta BF_10_ = 0.0005). Together, these results provide strong evidence against an effect of temporal presence on EEG power in the mu, theta, and beta frequency bands. See Supplementary materials S3 for analyses of neural responses to own pain.

### Exploratory analyses of whole time-frequency-electrode space

To analyze the neural effects of temporal presence and shock intensity beyond the hypothesized time windows and frequency bands, we conducted a whole-time-frequency-electrode space analysis (whole-brain analysis in the following) like we did in our previous study (Petereit et al., 2023). For this, we used permutation tests to create null hypotheses distributions for the effects of shock intensity, temporal presence condition, and their interaction. We compared our actual data with these null hypotheses data to identify significant EEG responses. Results were maximum value corrected. For the effect of shock intensity averaged over both conditions (Figure 4A), we found a significant cluster at the frequencies 1-6 Hz in the time window of 270-1450 ms after shock onset with maximum power at electrode Fpz. This indicates that observers’ frontal (low) theta activity is related to the observed pain of the target. A second significant cluster, showing a negative effect of shock intensity on power between frequencies 9-13 Hz, was found for the time window of 870-1550 ms after shock onset with the strongest deactivation at electrode PO8. This indicates that with increasing pain of the target, right posterior alpha activity in observers was reduced. For the effects of condition and the interaction of shock intensity and condition (Figure 4B-C), we found no activation that survived correction for multiple comparisons. Together, these exploratory whole-brain analyses revealed robust effects of shock intensity on frontal low-theta and posterior alpha activity, but no evidence for effects of temporal presence.

**Figure 4.**
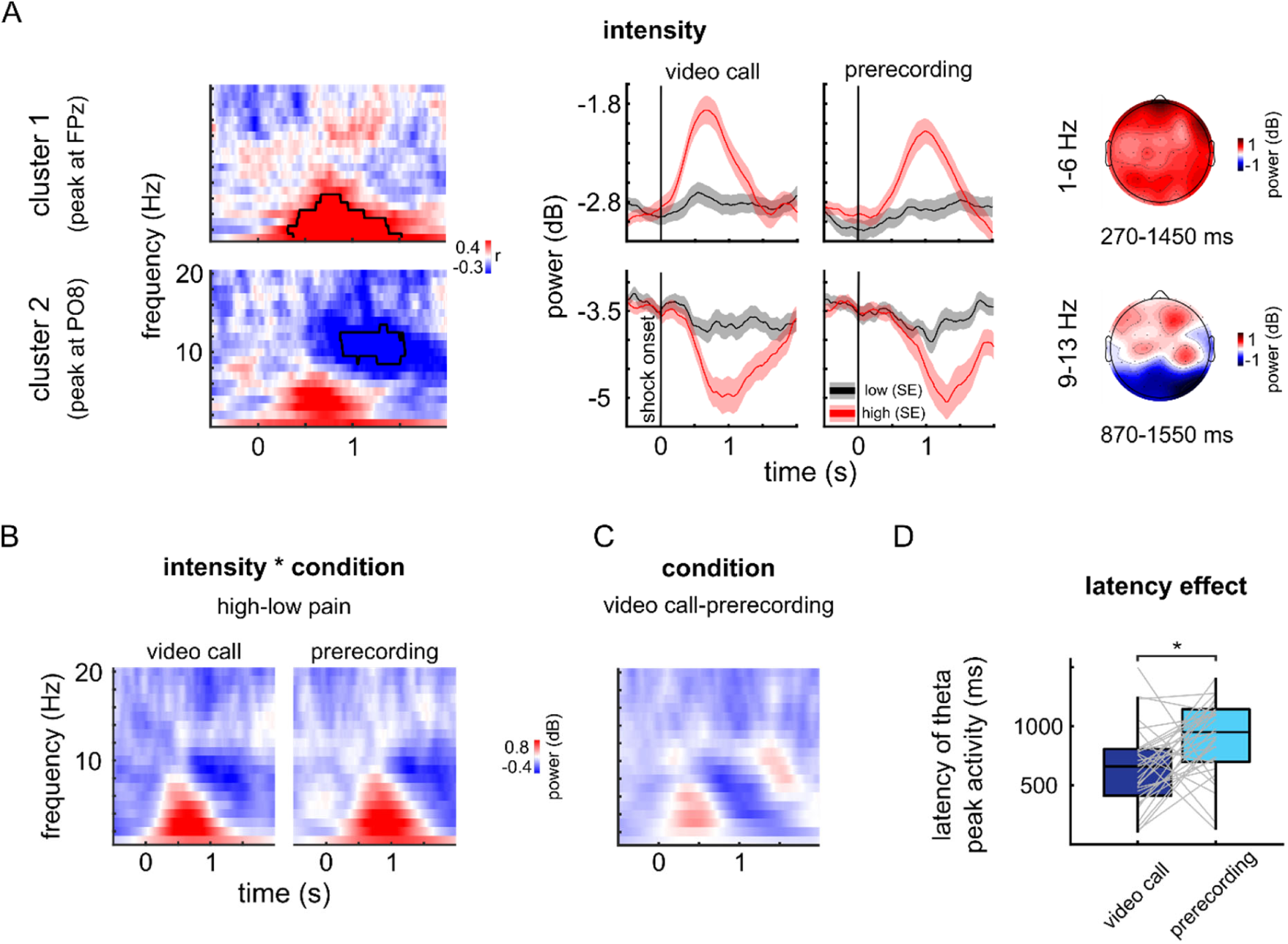
A) – C) Exploratory analysis of the whole time-frequency-electrode space. Shock intensities were dichotomized into low- and high-intensity for display purposes only. A) The effect of shock intensity. Two clusters survived the maximum-value correction. The time-frequency-power plots display correlations of power and pain intensities at electrodes of maximum amplitude for each cluster. Power time courses display the average power of the significant frequency range for each cluster. Topographies show the average power across the significant frequency and time range. B) The interaction of shock intensity and condition. Displayed is the effect of high-pain minus low-pain intensities per condition averaged across all electrodes. C) The effect of experimental condition: The heatmap shows the difference between conditions averaged across all electrodes and shock intensities. D) Exploratory latency analysis of the theta peak. Participants’ theta peak was delayed in the “prerecording” compared to the “video call” condition.

We used the results of the whole-brain analysis to explore the relationship between behavioral and neural empathic responses. On the single trial level, observer’s frontal theta activity (1-6 Hz, Fpz, 270-1450 ms) was weakly, positively correlated to observer’s pain ratings (“video call” condition: *r*(30) = .14, *p* < .001, *p_Bonferroni_* < .001; “prerecording” condition: *r*(30) = .13, *p* < .001, *p_Bonferroni_* < .001) and unpleasantness ratings (“video call” condition: *r*(30) = .13, *p* < .001, *p_Bonferroni_* < .001; “prerecording” condition: *r*(30) = .13, *p* < .001, *p_Bonferroni_* < .001), indicating a relationship between neural and behavioral empathic responses in empathy for pain.

### Exploratory analysis of the temporal presence effect on EEG activation latency

Our theta analyses were based on hypotheses drawn from our previous study. However, as can be seen in Figure 3A, the actual theta response of observers occurred later than expected (grey areas indicate hypothesized time windows). To investigate this, we conducted exploratory analyses on latency effects to determine whether temporal presence has an impact on peak latency in the theta and alpha bands. The frequency band ranges were based on the exploratory whole-brain analysis, which yielded effects for mid-frontal theta (1-6 Hz, Fpz) and right posterior alpha activity (9-13 Hz, PO8).

To test weather this latency effect was due to technical reasons and a delay of the video presentation in the “prerecording” condition, we analyzed the latency of the auditory N1 to the sound cue which indicated the shock onset. The auditory N1 peaked at 132 ms in the “video call” condition and at 141 ms in the “prerecording” condition, showing that there were no general delays in the “prerecording” condition (see Supplementary materials S4 for details).

As the theta and alpha activity delay was not caused by technical issues, we next tested whether it was statistically robust. We therefore identified the latencies of the maximum theta amplitude and of the minimum alpha amplitude on the individual subject level. We found a significant difference in theta peak latencies between conditions (Figure 4D; electrode Cz; *t*(30) = -2.78, *p* = .009, *p_Bonferroni_* = .019, *d* = 0.54). The average theta peak was at 649 ms in the “video call” and at 901 ms in the “prerecording” condition (difference: 252 ms). The analysis for the alpha decrease yielded no condition difference (electrode cluster: PO8; peak in the “video call” condition at 984 ms, peak in the “prerecording” condition at 1094 ms, *t*(30) = -1.37, *p* = .180, *p_Bonferroni_* = .360). These results indicate that there is a temporal presence effect on frontal theta activity with an earlier theta peak in the “video call” condition (higher temporal presence) compared to the “prerecording” condition (lower temporal presence). Notably, the observed latency effect of temporal presence was not only visible in the time-frequency data in the theta band, but also in the event-related potentials (see Supplementary materials S4).

We noticed that a similar effect might be evident in our previous study about the effects of physical presence on empathy for pain (Petereit et al., 2023). Therefore, we revisited the data from that study and found a similar latency effect (electrode Cz, latency of the theta peak in the “direct” condition: 636 ms; “mediated” condition: 808 ms; difference: 172 ms; *t*(28) = -1.92, *p* = .033, *d* = 0.30). This result also shows delayed latencies in the condition with reduced social presence. The time course of the theta power of the previous study can be seen in Figure 3D in Petereit et al. (2023). Thus, there seems to be a theta latency difference between social presence conditions in both studies.

### Temporal presence does not modulate physiological coupling between observers and targets

We used LMMs to test a possible effect of temporal presence on physiological coupling between observers’ and targets’ SCR and IBI. Therefore, we calculated whether observers’ data could be predicted by targets’ data and whether this effect was modulated by temporal presence. For both SCR and IBI, target’s data predicted the observer’s data, showing significant physiological coupling in terms of cardiac and electrodermal activity. However, no significant effect of condition or an interaction of target data and condition was found on physiological coupling. Model parameters are shown in Table 1. Testing the SCR model with and without the fixed effect of condition, the obtained Bayes factors of BF_10_ = 0.0009 indicated extreme evidence for the model without the predictor condition given our data. This indicates that the SCR coupling did not differ between the conditions with low and high temporal presence. Testing the IBI model with and without the fixed effect of condition, the obtained Bayes factors of BF_10_ = 0.0003 indicates extreme evidence for the model without the predictor condition given our data. This indicates that the IBI coupling did not differ between the conditions with low and high temporal presence either. Together, these results provide strong evidence that temporal presence does not modulate physiological coupling between observers and targets in either skin conductance responses or interbeat intervals. Observers’ SCR and IBI data in the observing conditions are shown in Figure 5A-B and physiological coupling is shown in Figures 5C-D. Neither SCR nor IBI physiological coupling were correlated to empathic accuracy (calculated across conditions; SCR: *r*(28) = .25, *p* = .178, *p_Bonferroni_* = .355; IBI: *r*(29) = .17, *p* = .362, *p_Bonferroni_* = .725).

**Figure 5.**
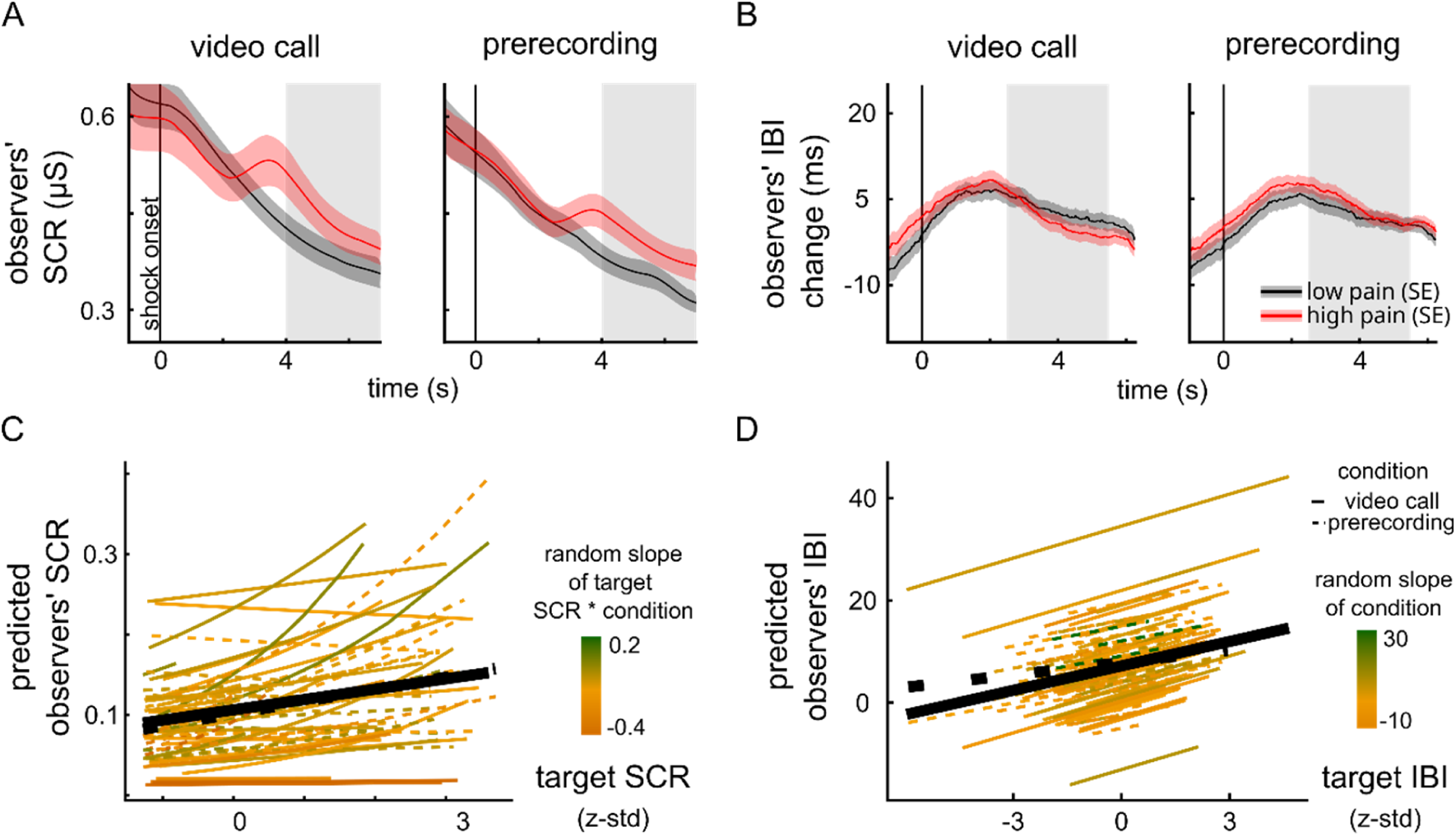
A) Observers’ SCR by condition. B) Observers’ IBI change by condition. Shock intensities were dichotomized into low- and high-intensity for display purposes only. Grey areas indicate time window of maximum correlation between target and observer data averaged across conditions. C) GLMM on skin-conductance coupling by predicting observers’ SCR with targets’ SCR. D) LMM on heartbeat coupling by predicting observers’ IBI with targets’ IBI. For both physiological read-outs, observers’ empathic responses were coupled to targets’ pain responses. However, this relationship was not affected by temporal presence. Black solid and dotted lines represent the “video call” and “prerecording” averages, respectively, while the green to yellow color coding reflects the strength of random slopes per participant.

### Condition order, habituation effects, and sex/gender effects

We did not find any condition order effects on observers’ pain and unpleasantness ratings, empathic accuracy, or affective empathy measures. Targets and observers showed habituation effects in their pain and (in case of observers) unpleasantness ratings. However, empathic accuracy and affective empathy did not change between first and second half of each condition. We did not find any effects of observers’ or targets’ sex/gender on empathic accuracy or affective empathy. Details can be found in Supplementary materials S5.

## Discussion

We examined pain empathy differences between a live, video-mediated interaction as opposed to watching a prerecorded video of a similar pain scenario. In both conditions, observers watched targets undergoing painful electric stimulation. Based on our theoretical framework, we expected reduced temporal presence to impair empathic processing. However, we found that observers were just as accurate in judging the other’s perception of pain when they passively watched a prerecorded video as when they were in a live video-mediated interaction. Moreover, observers experienced the other’s suffering as equally unpleasant in both conditions. At the neural level, we did not find mu suppression over the somatosensory cortex to be sensitive to the other’s pain in either condition. In contrast, mid-frontal theta did track the other’s pain intensity. Exploratory whole-time-frequency-space analyses confirmed this result and additionally suggested that a decrease in right posterior alpha power was involved in tracking the other’s pain intensity. However, these neural effects were independent of temporal presence. Further exploratory analyses revealed a possible latency effect of temporal presence on pain-related theta activation with delayed responses in the condition with reduced temporal presence. We observed significant skin-conductance and heartbeat coupling in the pain empathy paradigm, but this was also independent of temporal presence. In conclusion, the pain paradigm elicited meaningful empathic responses in observers at the behavioral, neural and physiological levels. However, these effects were not sensitive to temporal presence, with the exception of a latency effect of temporal presence on the theta response. The following section will first discuss the general findings regarding empathy for pain, and then discuss the null findings concerning the effects of temporal presence on empathy for pain.

### Behavioral, neural and physiological aspects of empathy for pain

In agreement with other studies on pain empathy, the observers in our study were able to accurately evaluate the other’s pain experience (Ben Adiva et al., 2024; Gauthier et al., 2008; Goldstein et al., 2017, 2018; Laursen et al., 2014; Leonard et al., 2013; Ruben & Hall, 2013; Shahri et al., 2024). The observers in our study showed high empathic accuracy, with distances to the targets’ ratings close to zero and a medium-sized correlation between observer and target pain ratings. Also consistent with other studies, participants showed affective empathy, reflected in increasing unpleasantness ratings in reaction to the other’s suffering (Fan & Han, 2008; Li et al., 2017; Mu et al., 2008; Peng et al., 2021; Petereit et al., 2023; Rütgen et al., 2015). Further, our findings are consistent with previous studies (Goldstein et al., 2017; Hein et al., 2011; Murata et al., 2020; Petereit et al., 2023; Reddan et al., 2020), which demonstrated that empathy for pain is associated with physiological coupling between the pain receiver and observer. Although the current study lacks a control condition without pain stimulation, Figure 5A shows that observers indeed exhibit an SCR when observing others in pain, indicating an empathic response. The relationship between physiological coupling and empathic accuracy seems to be less clear, since there are mixed results (Jospe et al., 2020; Levenson & Ruef, 1992; Thorson, 2018; Zerwas et al., 2021). Our results suggest no relationship between physiological coupling and empathic accuracy. Further research is therefore needed.

Regarding neural indices of empathy, we found that mid-frontal theta activity was related to others’ pain, replicating previous findings in pain empathy research (Misra et al., 2017; Mu et al., 2008; Peng et al., 2021; Petereit et al., 2023). Moreover, single-trial observers’ theta activity correlated with observers’ pain and unpleasantness ratings, demonstrating a close relationship between neural and behavioral indices of pain empathy. This correspondence was evident for both own pain (Supplementary Materials S3) and vicarious pain, supporting the interpretation that mid-frontal theta might reflect the perceived aversiveness or salience of others’ suffering (Legrain et al., 2011; Seeley, 2019; Uddin, 2015).

Contrary to our hypothesis and much of the previous literature (Li et al., 2017; Peled-Avron et al., 2018; Perry et al., 2010; Riecansky & Lamm, 2019; Whitmarsh et al., 2011), we did not observe mu suppression in response to observing others’ pain. As we did observe clear mu suppression in the “own pain” condition, and as we applied a Laplace transformation (current source density, CSD) to improve separation of occipital alpha and sensorimotor mu, we are confident that this absence of mu suppression is not a methodological artefact. Notably, the null finding replicates our earlier study using the same pain empathy paradigm (Petereit et al., 2023), suggesting that it is not incidental but may be characteristic of this specific experimental context. One possible explanation is that mu suppression is particularly sensitive to the observation of overt actions or somatosensory events (Fox et al., 2016; Perry & Bentin, 2009; Whitmarsh et al., 2011), whereas the present paradigm primarily relied on facial expressions of pain without visible limb movement or direct action observation. In such contexts, affective and evaluative components of empathy may be more strongly engaged than sensorimotor mirroring mechanisms (Saarela et al., 2006). Importantly, the absence of mu suppression occurred alongside robust behavioral, physiological, and midfrontal theta responses to others’ pain, indicating that empathic processing was preserved despite the lack of a sensorimotor mirroring signature. Taken together, these findings add to concerns about the reliability and specificity of mu suppression as a general neural marker of empathy (Hobson & Bishop, 2016) and suggest that its involvement may depend on task characteristics and the type of empathic cues available.

### Effect of temporal presence on empathy for pain

We did not find effects of temporal presence on behavioral indices of pain empathy. Moreover, we did not find temporal presence effects on neural mu, beta and theta activity and also not on physiological markers of empathy for pain. However, there was a tendency of reduced theta power in the condition without synchronous exchange of information compared to the synchronous condition (see Figure 4A). Thus, with more power this effect might have reached significance. Furthermore, we identified a temporal presence latency effect on the theta peak, which indicates delayed responses without synchronous information exchange.

The results of the current study are similar to our previous study, which compared direct, face-2-face and video call interaction (“physical presence”). In this study (Petereit et al., 2023), we did not find effects of physical presence on behavioral indices of pain empathy, mu and beta power, and heartbeat coupling. But in contrast to the current study on temporal presence, there were effects of physical presence on theta power and EDA coupling in relation to pain empathy. This suggests that physical presence, i.e., the number of available social cues, might have a stronger influence on the responses to others’ pain than the synchronous exchange of information between pain receiver and observer. This demonstrates that the empathic physiological coupling between individuals is preserved without bidirectional exchange of social cues when comparing live video calls and viewing prerecorded videos.

Diana and colleagues (2023) found mimicry and trust to be lower in response to prerecorded videos compared to direct, live interaction, which suggests that experience-sharing might be diminished with reduced temporal presence. In contrast, we found that empathic responses to others’ pain were not affected by temporal presence. This suggests that our empathic abilities, like the understanding and sharing of other’s pain, are not impaired by lacking temporal presence. This is in line with Morgan and colleagues (2022), who found that explicit mentalizing was unaffected by social presence when comparing direct, live interactions and prerecorded videos in a theory of mind task.

As far as our measures of empathy are concerned, our results suggest that temporal presence, and thus the possibility of reciprocal exchange (at least non-verbally in this case), does not modulate a person’s empathy for the other. Schilbach and colleagues (2013) stressed “the importance of experiencing and interacting with others as our primary ways of knowing them” (p. 395). In this sense, the emotional engagement and the social interaction with its complex interaction dynamics between agents are considered key constituents of social cognition. Similarly, De Jaegher and colleagues (2010) emphasized that co-regulation of coupling between interacting individuals is crucial for social understanding. The relational dynamics and connectedness between individuals are supposed to affect the cognitive processes involved in thinking about the other. Provided that our measures for empathy are adequate, our results suggest that this is not necessarily the case. In our paradigm, the participants were able to understand, share the other’s pain, and physiologically resonate with the other just as much without synchronous interaction and by merely observing the other’s pain expression. Our data suggest that the pain perceivers were as emotionally engaged (measured as affective empathy) in a non-interactive condition as they were in an interactive condition. Thus, for the domain of pain empathy, understanding and sharing the other’s state might not be affected by the temporal presence between pain receiver and observer. Empathy for pain likely relies on less complex social cues, which enables understanding and sharing of experiences even without synchronous exchange of cues between observer and target. This may contrast with more complex social interactions, such as sharing negative autobiographical emotional events (Jospe et al., 2020), which may depend more heavily on feedback loops of social exchange.

It can be said, however, that even the live, video mediated condition did not really offer opportunities for social interaction. The participant’s task was to estimate the other’s pain intensity and rate their own unpleasantness in response to the other’s suffering, but they were not asked to interact or respond to the other, or to display prosocial behavior. In contrast, due to the constraints of the EEG recordings, they were asked to refrain from movements and talking. A paradigm that requires and allows for more interaction might therefore show more pronounced effects of temporal presence on empathy. For instance, more complex socio-cognitive phenomena, such as cooperation, prosocial motivation, and trust, may be more sensitive to temporal presence. In comparison, empathy for pain may rely on relatively direct, perceptual, affective cues. Nevertheless, our results are informative for second-person neuroscience (Cañigueral et al., 2022; De Jaegher et al., 2010; Gallotti et al., 2017; Schilbach et al., 2013), as they show that at least when it comes to empathic responses to others’ pain, they remain preserved and unaltered also with reduced temporal presence and reciprocal interaction between individuals. Put more positively, our results are reassuring for social neuroscience research using videos as stimuli, because the differences to direct interactions might not be as strong as one might expect.

We cannot exclude though that temporal presence has an effect on empathy beyond those measures of empathy we assessed here. Not being able to interact with the other person due to reduced temporal presence may have reduced the self-relevance of the other person (Conway et al., 2016; Schmitz & Johnson, 2007) and thereby the motivation for prosocial behavior or for showing compassion (Ferguson et al., 2020; Hein et al., 2011; Weisz & Zaki, 2018; Zaki, 2014). As our experimental setup did not allow the observers to express their compassion – other than through facial expressions – and as we did not assess their motivation for prosocial behavior, this remains speculative. Future social presence research should consider measuring motivational aspects of empathy.

Although the results of our hypothesis testing indicated no effect of temporal presence on empathy for pain, we did identify a latency effect of temporal presence on frontal theta activity. Participants’ frontal theta activity in response to the others’ pain was delayed in the condition with reduced temporal presence with a difference of about 252 ms. This latency shift constitutes the primary neural marker of reduced temporal presence in the present study, indicating an alteration in the temporal dynamics of empathic processing rather than its overall magnitude. Possibly, differences in visual attention and self-relevance could explain the latency effect. On a behavioral level, self-relevance has been found to speed up visual search responses (Wade & Vickery, 2018) and to impact visual attention (Svensson et al., 2023). On the neural level, self-relevant information increases neural responses (Sui et al., 2013; Viskontas et al., 2009). Not being seen by the observed person may lead to differences in attending visually to the others’ pain expression compared to social encounters in which both participants can see each other (Hietanen et al., 2020; Morgan et al., 2022). Thus, differences in self-relevance may have affected attention processes in the current study, reflected in delayed neural responses.

Interestingly, revisiting the data from our previous study about the effects of physical presence on empathy for pain revealed a similar latency effect. The latency of the theta peak in the condition with reduced social cues (live video call) was 172 ms later than in the condition with rich social cues (direct, face-to-face). Together, these findings suggest that reduced social presence - across both temporal and physical dimensions - consistently affects the timing of neural responses associated with empathy for pain. Further studies are needed to specify the cognitive or motivational factors that are relevant here, be it attention and self-relevance or other factors. The combination of preserved behavioral responses and physiological coupling in empathy for pain with delayed frontal theta activity under reduced temporal presence may indicate that empathic processing remains largely intact, while its neural timing is altered. Rather than reflecting a reduction in empathy, the latency shift could point to subtle changes or compensatory mechanisms in the processing dynamics supporting empathic understanding when temporal presence is reduced.

### Limitations

As with any null effect, the lack of condition differences in the current study could be due to the sample size and too limited power. However, the prior power analysis suggested that a sample of thirty participants is sufficient to detect also a small behavioral effect in a within-subject design. Moreover, the Bayes factor analyses indicated quite strong evidence for the null hypotheses.

Our sample was quite homogeneous, consisting of young, predominantly female students. This limits generalizability to male, older and non-student populations. In addition, younger people may be more adept at digitally mediated interactions (Vossen & Valkenburg, 2016), which could offset any impacts of temporal presence on empathy. Vice versa, effects of temporal presence on empathy might be more pronounced in populations with less experience in digitally mediated social interactions.

Furthermore, the homogeneity of the sample might have contributed to the generally high levels of empathic accuracy, as previous research has shown that people are more empathically accurate with similar others (Han, 2018; Majdandžić et al., 2016) and with familiar others (Israelashvili & Perry, 2021). We only included dyads of strangers and guaranteed low levels of familiarity. However, as participants were all similarly young students of the same University, it is likely that they perceived each other as quite similar. The fact that participants were not more accurate in the live video calls compared to the prerecorded video condition may be thus due to a ceiling effect. Future studies on social presence might therefore consider including more heterogenous samples.

Although the “video call” condition met the criteria for an actual video call, the participants remained in close proximity, and knew they were in adjacent rooms. Due to the within-subject design of the study, it was not possible to physically separate the participants further. This limits the generalizability of our findings, since real video calls usually also involve greater physical distance, and the person’s knowledge about this greater physical distance might lower the relevance of the other person for the self. Future studies will have to show whether, in addition to technical transmission, the actual physical distance plays a role for the participants’ behavioral, physiological, and neural empathic responses.

In terms of technical limitations, it should be noted that the hand, to which EDA electrodes were attached, was also used for operating the joystick. Unfortunately, operating the joystick for the VAS ratings introduced artifacts to the EDA data, which made it necessary to exclude trials. With more trials and therefore more power, we might have been able to discover the probably nuanced effects of temporal presence in the EDA data, as there was at least a trend for a condition difference (see Figure 5A).

## Conclusion

Digitally-mediated social interactions have become increasingly common in daily life. Research has just begun to explicitly compare direct and mediated interactions (Diana et al., 2023; Hietanen et al., 2020; Järveläinen et al., 2001; Karimova et al., 2024; Prinsen & Alaerts, 2019; Shimada & Hiraki, 2006), especially with regard to social cognition (Jospe et al., 2020; Morgan et al., 2022; Petereit et al., 2023; Pönkänen et al., 2011). In the current study on temporal presence and pain empathy, we found no evidence that a lack of temporal presence between individuals significantly affects empathy for pain. This indicates that major aspects of empathic responses to other’s pain are unaltered in mediated interactions. At the neural level, reduced temporal presence was primarily reflected in a latency shift of frontal theta activity, suggesting an alteration in the temporal dynamics of empathic processing rather than in its overall magnitude. Therefore, our findings are relevant for second-person neuroscience (Schilbach et al., 2013), but also for online psychotherapy and telemedicine (Amichai-Hamburger et al., 2014; Grondin et al., 2019). If empathic responses to others’ pain are preserved when watching a prerecorded video, then the cost of online therapy through live video interaction might be negligible, at least in terms of understanding others’ pain. Future studies will need to examine the effects of social presence on empathy in more complex contexts and considering also motivational aspects of empathy.

## Data and Code availability

The raw data, the final preprocessed data, which is prepared for statistical analysis, and the analysis code can be found at https://osf.io/3jdf4/ (DOI 10.17605/OSF.IO/3JDF4).

## Acknowledgements

We would like to thank Lea Henke, Kira Weckmann, Liina Elsner, and Nadine Horchler for their assistance in acquiring the data. We also thank Andreas Sprenger and Marcus Heldmann for their technical support.

## AI Usage Statement

Artificial intelligence tools (ChatGPT and DeepL) were employed only to assist with language editing and sentence-level rephrasing. All scientific concepts, analyses, and conclusions are entirely the author’s own or based on the previous scientific literature.

