## Supplementary materials for "Empathy for pain persists across live two-way video interactions and viewing of prerecorded videos"

#### S1: Information about questionnaires that have been assessed but not analyzed

After each experimental session, subjective measures, including discomfort, sympathy, and attractiveness, were assessed. However, these served as auxiliary information and were excluded from subsequent analyses since these variables are not directly linked to the research question regarding the impact of temporal presence on empathy. Following the experiment, participants completed a series of questions regarding their age, self-identified sex/gender, body weight, height, university degree, and habits concerning physical activity, smoking, and caffeine intake. Lastly, participants completed the Interpersonal Reactivity Index (Davis, 1983; German version: Paulus, 2009), which is also not reported here.

#### S2: R code for the linear mixed models and explanation of the variables

The dependent variables are the observers' pain ratings, unpleasantness ratings, EEG power in each frequency band, observers' skin conductance and observers' inter-beat interval data. The independent variables are the condition ("video call" vs. "prerecording") in interaction with the targets' pain ratings, the shock intensity, targets' skin conductance or targets' inter-beat interval data. The R code for each model is reported below.

1. Empathic accuracy (lmer function):
   obs_rating ~ 1 + cond * targ_rating + (1 + cond * targ_rating | subj_id)
2. Affective empathy (lmer function):
   unpleasantness ~ 1 + cond * targ_rating + (1 + cond * targ_rating | subj_id)
3. Observers' theta activity (lmer function):
   EEG_power ~ 1 + cond * SI + (1 + cond + SI | subj_id)
4. Observers' mu activity (lmer function):
   EEG_power ~ 1 + cond * SI + (1 + cond | subj_id)
5. Observers' beta activity (lmer function):

EEG_power ~ 1 + cond * SI + (1 | subj_id)

6. SCR coupling (glmer function):

   EDA_obs ~ 1 + cond * EDA_targ + (1 + cond * EDA_targ | subj_id)

7. IBI coupling (lmer function):

   IBI_obs ~ 1 + cond * IBI_targ + (1 + cond | subj_id)

Legend:

| | |
|---|---|
| cond | = condition (“video call” vs. “prerecording”) |
| obs_rating / targ_rating | = observers’ or targets’ pain ratings |
| unpleasantness | = observers’ unpleasantness ratings |
| EEG_power | = power of each frequency band in the observers’ EEG data |
| EDA_obs / EDA_targ | = observers’ or targets’ skin conductance data |
| IBI_obs / IBI_targ | = observers’ or targets’ inter-beat interval data |
| SI | = shock intensity |
| subj_id | = subject identity |

**S3: Behavioral and neural responses to own pain**

For EEG data of the “own pain” condition, we calculated LMMs for mu, theta, and beta activity. The shock intensity served as fixed effect. The best models included random effects for the intercept and the shock intensity. For mu, we analyzed electrode C3 at 8-12 Hz in the time window of 500-1000 ms (Genzer et al., 2022; Hoenen et al., 2015; Perry et al., 2010). For theta, we analyzed electrode Fz at 3-9 Hz in the time window of 0-500 ms (Misra et al., 2017; Ploner et al., 2017). For beta, we analyzed electrode CP1 at 11-20 Hz in the time window of 500-1000 ms (Petereit et al., 2023).

The LMM on behavioral data showed that with increased shock intensity, participants reported higher own pain ratings. For the neural responses, the LMM on participants’ mu activity indicated stronger mu suppression with increased shock intensity. Further, an LMM on theta activity indicated that theta power was stronger in reaction to increased painful stimulation. Finally, the LMM on participants’ (low) beta activity indicated stronger beta suppression with increased shock intensity. These results demonstrate that mu suppression, theta, and (low) beta activity are involved in pain processing (see left column of Figure 3A-C). Model parameters can be found in Table S1. On the single trial level, self-reported pain ratings were weakly correlated with both mu suppression ($r(27) = -0.189$, p < 0.001) and theta increase (r(27) = 0.147, p < 0.001), indicating a direct link between mu and theta responses to pain and subjective pain experience. Beta activity was not correlated with self-reported pain ratings (r(27) = -0.018, p < 0.001).

**Table S1** Results of the single-trial linear mixed models on behavioral, mu, theta, and beta data for own pain.

| Model | Random effects | SD | Fixed effects | b (SE) | t (df) | p | $R^2$ marg<br>$R^2$ cond |
|---|---|---|---|---|---|---|---|
| Targets' pain ratings (n = 31) | Intercept | 8.5 | Intercept | 39.83 (1.55) | 25.6 (30) | <0.001 | 0.675 |
| | Intensity | 5.6 | **Intensity** | 24.03 (1.03) | 23.3 (30) | **<0.001** | 0.796 |
| Targets' mu power (n = 30) | Intercept | 1.98 | Intercept | -5.5 (0.37) | -14.9 (29) | <0.001 | 0.04 |
| | Intensity | 0.6 | **Intensity** | -0.74 (0.13) | -5.57 (31) | **<0.001** | 0.32 |
| Targets' theta power (n = 30) | Intercept | 0.94 | Intercept | 0.2 (0.18) | 1.1 (28) | 0.28 | 0.012 |
| | Intensity | 0.32 | **Intensity** | 0.32 (0.09) | 3.69 (28) | **<0.001** | 0.137 |
| Targets' beta power (n = 30) | Intercept | 1.24 | Intercept | -3.82 (0.23) | -16.4 (29) | <0.001 | 0.007 |
| | Intensity | 0.32 | **Intensity** | -0.23 (0.08) | -2.85 (30) | **0.008** | 0.23 |

Notes: IBI = interbeat interval, SCR = skin conductance response, Intensity = shock intensity, $R^2$ marg = $R^2$ marginal (explained variance with fixed effects model), $R^2$ cond = $R^2$ conditional (explained variance with fixed and random effects model). Models on behavioral data, EEG power, and IBI responses are linear mixed models, and models on SCRs are generalized linear mixed models (note that the former provide Satterthwaite's degrees of freedom and that the latter do not provide degrees of freedom).

**S4: Auditory N1 analysis and pain empathy event-related potential**

To verify that we did not have any general delay between the video presentation in the "prerecording" and the "video call" condition, we analyzed the latency of the auditory N1 to the sound cue that indicated the shock onset at electrode Cz (Woods, 1995). A clean N1 could be obtained for 29 subjects. The auditory N1 peaked at 132 ms in the "video call" condition and at 141 ms in the "prerecording" condition (t(28) = -3.56, p = 0.001). Although being significant, the difference of 9 ms (SD = 13.5 ms) cannot explain the the difference in peak theta latency (252 ms) between conditions, which was 28 times greater than the N1 latency discrepancy (see Figure S1A).

In the main article, we described a theta latency effect of temporal presence (section "Exploratory analysis of the temporal presence effect on EEG activation latency"). This latency effect can also be seen in the pain-empathy event-related potentials (Figure S1B), which adds support for the latency effect of temporal presence beyond the theta frequency range. The topographies (Figure S1C) show the late positive potentials related to pain empathy, which has been demonstrated before (Mella et al., 2012).

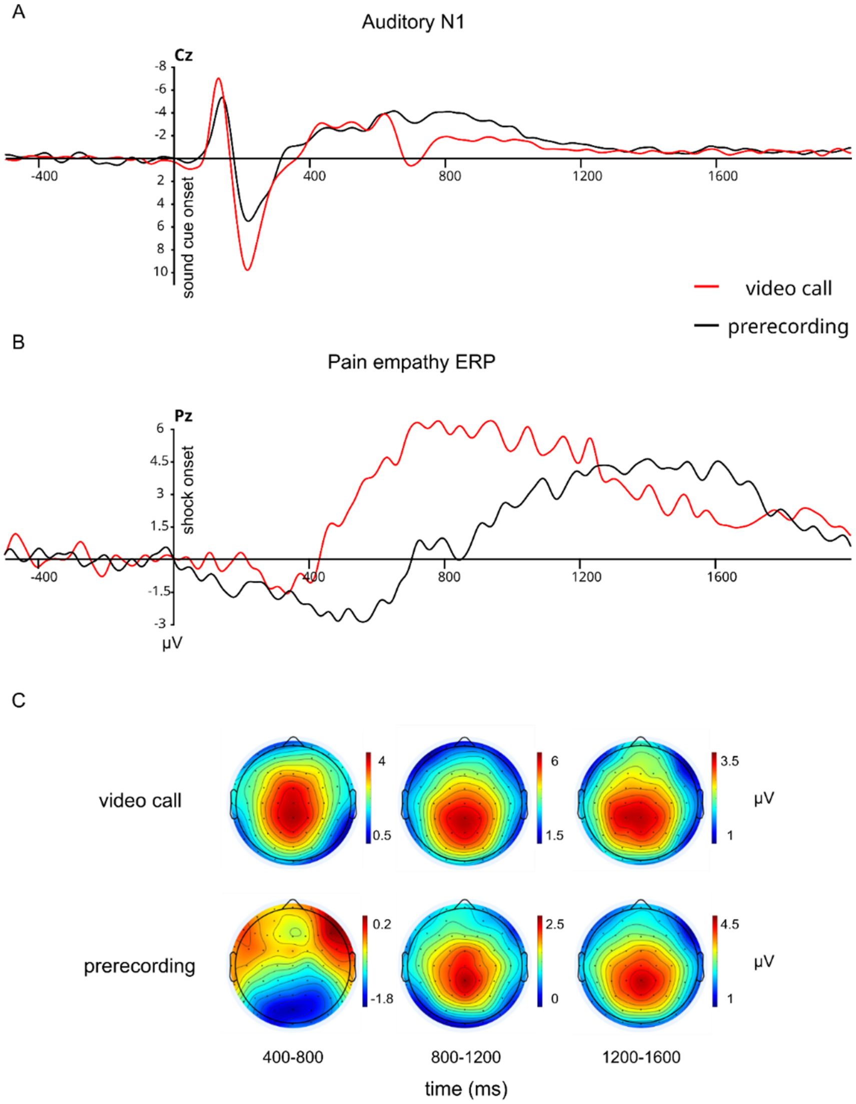


**Figure S1:** A) Auditory ERP for the "video call" and "prerecording" condition after the cue sound before the shock onset. The difference of 9 ms between N1 peaks indicates that the theta latency differences between the conditions were not caused by technical issues. B) Empathy-for-pain ERP for the difference of high-pain minus low-pain per condition, which illustrates the latency delay in the "prerecording" condition. C) ERP-topographies for the difference of high-pain minus low-pain per condition.

### S5: Condition order, habituation effects, and sex/gender effects

#### *No condition order effects*

To control for condition order effects, we tested for various measures if there were differences between first and second observing condition for each subject. The tested measures were the maximum pain stimulus calibration (first calibration before "own pain", second before "video call" condition), observers' pain ratings, targets' pain ratings and observers' unpleasantness ratings, as well as empathic accuracy and affective empathy. A paired t-test was used for testing of condition order effects in maximum pain stimulus calibration. For all behavioral rating analyses, LMMs were used similar to the behavioral data analysis but with an added dichotomous condition order predictor (video call -> prerecording vs. prerecording -> video call).

We did not find any differences between the first and second pain stimulus calibration ($t(33) = 0.38$, $p = 0.7$). Also, there were no condition order effects on observers' pain ratings (independent variable: condition order; dependent variable: observer pain rating; $b = -5.32$, $SE = 4.49$, $t(29) = -1.18$, $p = .247$), targets' pain ratings (independent variable: condition order; dependent variable: target pain rating; $b = 0.07$, $SE = 0.11$, $t(29) = 0.69$, $p = .494$), observers' unpleasantness ratings (independent variable: condition order; dependent variable: observer unpleasantness rating; $b = 0.55$, $SE = 6.75$, $t(29) = 0.08$, $p = .936$), empathic accuracy (independent variables: condition × target pain rating × condition order; dependent variable: observer pain rating; $b = -0.47$, $SE = 2.94$, $t(27) = -0.16$, $p = .875$), or affective empathy (independent variables: condition × target pain rating × condition order; dependent variable: observer unpleasantness rating; $b = -0.38$, $SE = 2.41$, $t(28) = -0.16$, $p = .874$). Therefore, we conclude that there were no ordering effects and no learning effects that improved participants' empathic abilities from first to second condition.

#### *Habituation in individual ratings but no change in empathic accuracy and affective empathy*

To control for habituation effects, we tested if there are differences in ratings between the first and second half within each condition. The same procedure was used as for condition order effects but with a dichotomous predictor for first and second half of each condition.

We found habituation effects for individual pain ratings of observers (independent variable: first vs. second half; dependent variable: observer pain rating; b = -2.04, SE = 0.94, t(2448) = -2.17, p = .030; mean rating first half: 41.1, second half: 39.0), pain ratings of targets (independent variable: first vs. second half; dependent variable: target pain rating; b = -0.19, SE = 0.04, t(2447) = -4.82, p < .001; mean rating first half: 42.5, second half: 37.1) and unpleasantness ratings (independent variable: first vs. second half; dependent variable: observer unpleasantness rating; b = -3.11, SE = 0.76, t(2422) = -4.11, p < .001; mean rating first half: 31.0, second half: 28.0). However, we did not find any changes in empathic accuracy (independent variables: condition × target pain rating × condition order; dependent variable: observer pain rating; b = -1.64, SE = 1.42, t(2364) = -1.16, p = .248), and affective empathy (independent variables: condition × target pain rating × condition order; dependent variable: observer unpleasantness rating; b = -1.48, SE = 1.18, t(2337) = -1.26, p = .209) within each condition.

#### ***No sex/gender effects on empathic accuracy or affective empathy***

We examined for potential sex/gender differences in empathic accuracy and affective empathy by examining interactions involving observer sex and target sex. Specifically, we assessed the three-way interactions between temporal presence condition, pain rating, and sex/gender (observer or target) using separate linear mixed models. Our sample consisted of 28 females and seven males (self-identified).

For empathic accuracy, the three-way interaction with observer sex (independent variables: condition × target pain rating × observer sex/gender; dependent variable: observer pain rating) was not significant (b = 2.19, SE = 1.81, t(2264) = 1.21, p = .227), nor was the interaction with target sex (independent variables: condition × target pain rating × target sex/gender; dependent variable: observer pain rating; b = 0.12, SE = 1.83, t(2411) = 0.06, p = .949).

Similarly, in the affective empathy models, the three-way interaction including observer sex was non-significant (independent variables: condition × target pain rating × observer sex/gender; dependent variable: observer unpleasantness rating; b = 3.98, SE = 2.88, t(28) = 1.38, p = .178), as was the interaction with target sex (independent variables: condition × target pain rating × target sex/gender; dependent variable: observer unpleasantness rating; b = -0.17, SE = 3.02, t(27) = -0.06, p = .955).

Overall, there were no statistically significant three-way interactions involving sex/gender for either empathic accuracy or affective empathy, indicating that sex of the observer or target did not modulate the relationship between condition and target ratings on empathy measures.
